# A large, comprehensive labeled MRI dataset of the human lateral ventricle choroid plexus

**DOI:** 10.64898/2026.10.08.757728

**Authors:** Ling-Yu Huang, Victor Zeng, Nicolas Raymond, Brendan Stiltner, Willa Molho, Yuan Cao, Deepthi Bannai, Daphne Ying, Emma Oss, Daman P. Dhunna, Prachi Patel, Alexa Adesokan, Morgan Hricay, Victoria Lyons, Noah Tolentino, Matcheri S. Keshavan, Peter Fried, Mohammad Jammoul, David Kuegler, Santiago Estrada, Martin Reuter, Paulo Lizano

**Affiliations:** Department of Psychiatry, University of Rochester Medical Center, Rochester, NY, USA; Department of Psychiatry, Beth Israel Deaconess Medical Center, Boston, MA, USA; Department of Psychiatry and Psychotherapy, Jena University Hospital, Jena, Germany; German Center for Mental Health Halle-Jena-Magdeburg Site, Jena, Germany; Redwood Center for Theoretical Neuroscience, University of California Berkeley, Berkeley, CA, USA; German Center for Neurodegenerative Diseases, Bonn, Germany; Department of Radiology, Harvard Medical School, Boston, MA, USA; A. A. Martinos Center for Biomedical Imaging, Massachusetts General Hospital, Charlestown, MA, USA

## Abstract

The choroid plexus (ChP) is a highly vascularized structure located in the brain ventricles that plays important roles in multiple central nervous system functions. ChP enlargement has been implicated in many neuropsychiatric conditions such as multiple sclerosis, depression, schizophrenia, and Alzheimer’s disease. However, limitation in the accuracy of the current segmentation tools highlight the need for more reliable methods to quantify ChP volume in healthy and clinical human populations. The shape and size of the ChP vary drastically from person to person, and currently there are no large ChP datasets which allow comprehensive training of segmentation models. Here we introduce DataPlex, a collection of 700 structural MRI brain images where the lateral ventricle ChP has been manually labeled and checked by expert human raters. DataPlex is balanced and representative across a wide range of scanning parameters, demographics, and clinical profiles, and subject-level information will be available for each labeled image. DataPlex will be open source to support the creation of reliable segmentation tools for the human lateral ventricle ChP.

## Introduction

The choroid plexus (ChP) is a specialized tissue found in each of the four ventricles of the brain (**Figure 1a**). ChP consists of a monolayer of epithelial cells surrounding fenestrated capillaries and forms the blood-CSF barrier (B-CSF-B) while producing cerebrospinal fluid (CSF), thereby contributing to brain homeostasis, neurodevelopment, immune surveillance, and waste clearance through the glymphatic system^1–5^. Compared to other brain regions, the understanding of ChP pathophysiology differs across neuropsychiatric disorders, including its morphology, and the literature is relatively sparse^6^. Although the ChP has long been recognized for its physiological functions, its role in neuropsychiatric disorders remains poorly understood^6^. Increasing evidence has reported enlarged lateral ventricle ChP across multiple neuropsychiatric and neurodegenerative disorders, including psychosis spectrum disorders, depression, bipolar disorder, and Alzheimer’s^7–14^.

**Figure 1.**
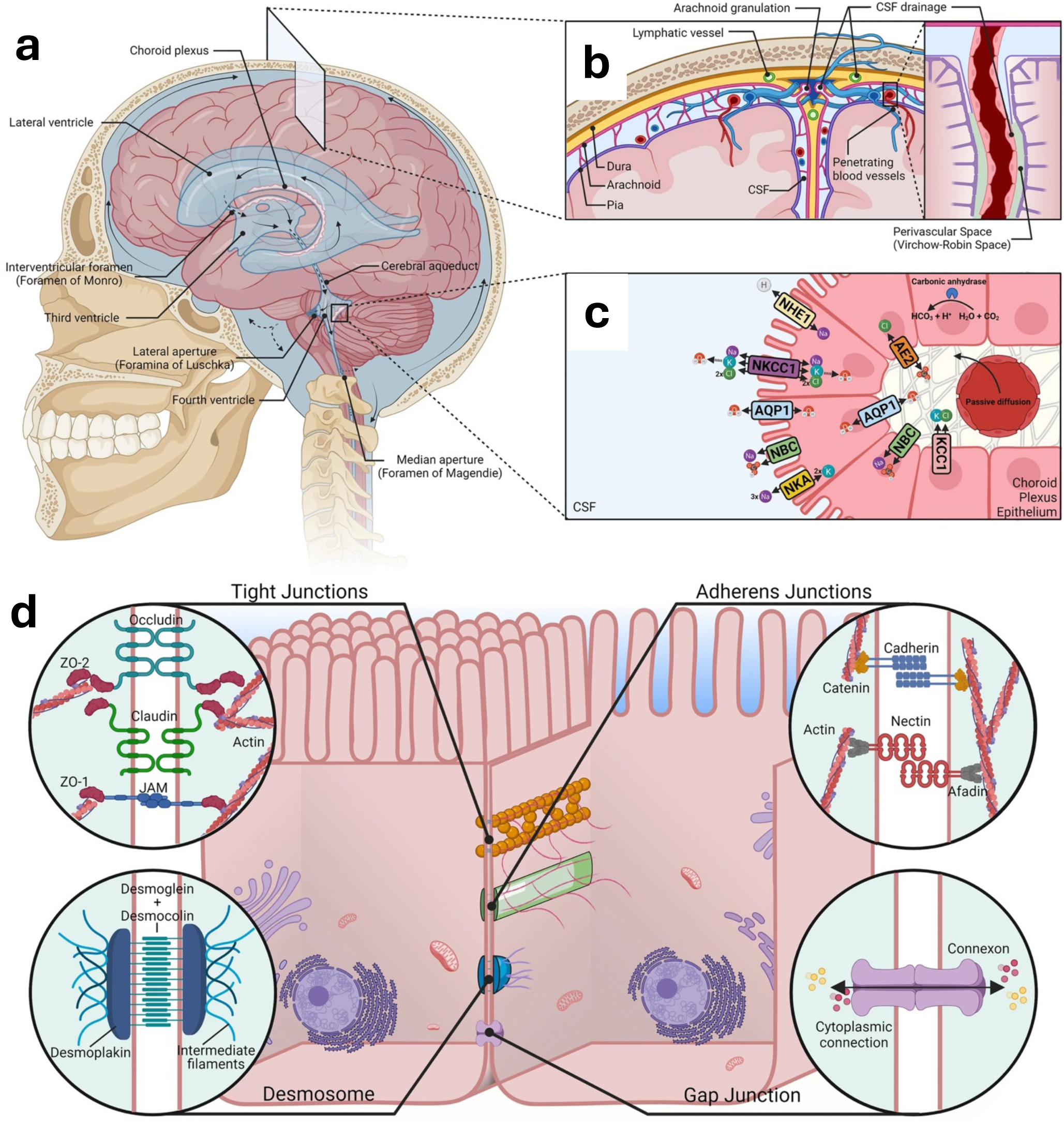
Overview of the role of the ChP as the blood-CSF barrier and driver of glymphatic movement. **a.** The arrows indicate the direction of CSF flow. Pressure created by CSF production at the ChP causes the fluid to flow unidirectionally within the ventricles. The flow becomes multidirectional after exiting through the Foramen of Luschka (lateral aperture) and and Foramen of Magendie (medial sperture) near the fourth ventricle. **b.** A cross-sectional view showing the three drainage routes of CSF: lymphatic tunnels, arachnoid granulation, and perivascular spaces. Flow of CSF through the perivascular spaces is especially important for glymphatic cleansing of metabolic waste from brain tissues. **c.** Morphological and molecular asymmetry exists and is maintained across the apical and basal sides of the ChP epithelial membrane. Expression of ion pumps and co-transporters determine the membrane potential and concentration gradients, which in tern determine the direction of movement for solutes and water molecules. The asymmetry in membrane protein expression allows control of CSF production and other long-range signaling effects. **d.** Tight junctions, adherents, and other intracellular anchors ensure that chemical traffic entering or exiting the CSF is constrained to the regulation of ChP membrane channels instead of free-flowing through the paracellular spaces. (adapted from Thompson et al. (2022)^37^)

Structural magnetic resonance imaging (MRI) provides a non-invasive approach for measuring ChP morphology in-vivo and can be readily applied across large cross-sectional and longitudinal studies. Because the ChP can be distinguished from the surrounding cerebrospinal fluid on high-resolution T1-weighted MRI (**Figures 2, 3, S1, and S2**), MRI has become the primary imaging modality for studying ChP volume in humans. However, accurate quantification remains challenging. Manual segmentation is currently considered the gold standard but typically requires 30-60 minutes to complete per scan, making it impractical for large imaging cohorts. In addition, manual delineation introduces inter-rater variability, and there is a shortage for standard labeling protocols and reliable anatomical labels, particularly across datasets acquired using different scanners, manufacturers, and imaging protocols. These limitations highlight the need for accurate, reproducible, and automated methods for ChP segmentation.

**Figure 2.**
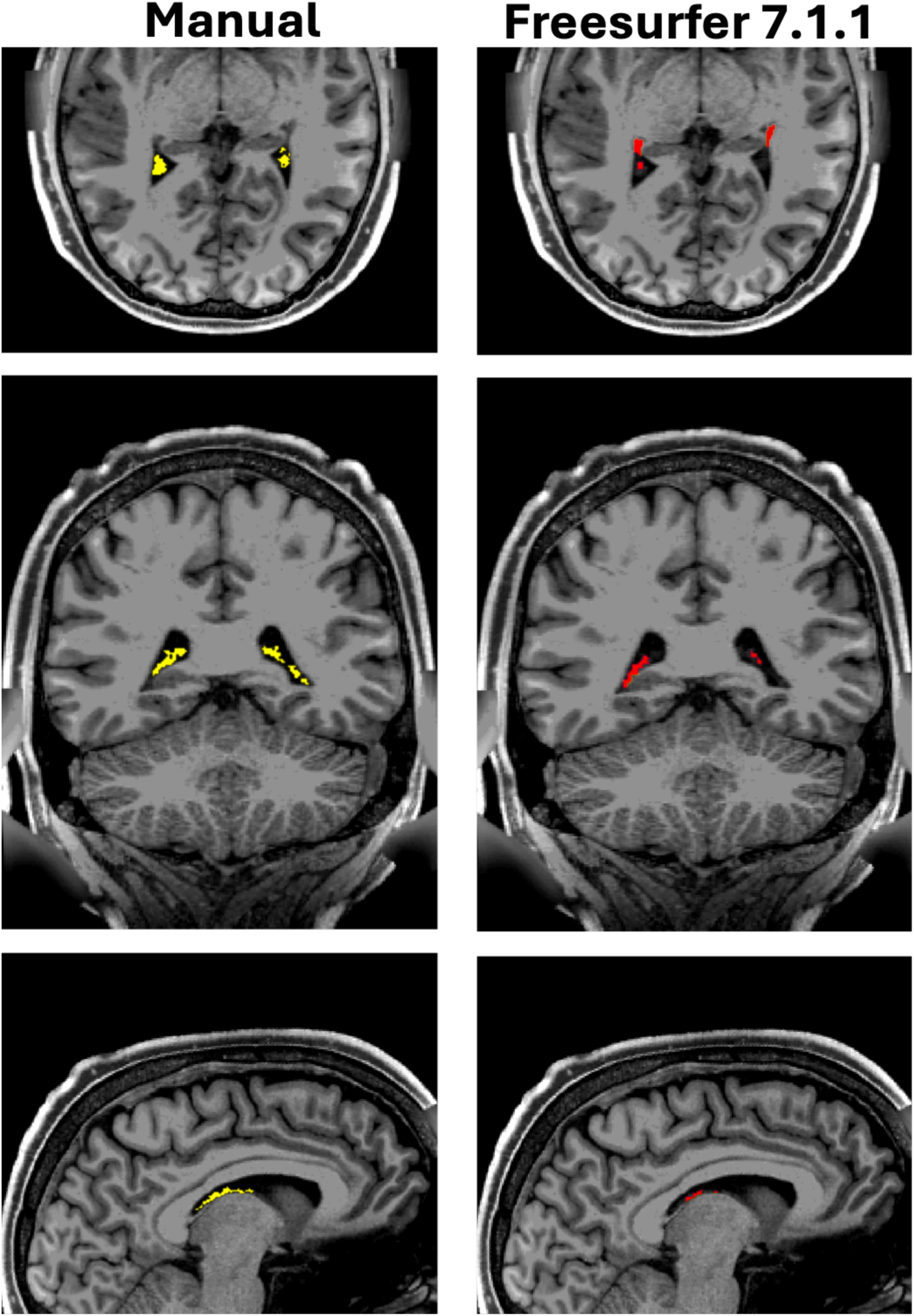
Comparison of segmentation quality between FreeSurfer 7.1.1, the only open-source analysis software for human structural MRI segmentation of the lateral ventricle ChP, and manual segmentation by an expert human rater. FreeSurfer performed poorly in both placement and volume estimation. Manual segmentation is currently the “gold standard” for estimating the volume of the human lateral ventricle ChP, but it is time-consuming and therefore not realistic for large imaging consortia with thousands of participants.

A limited number of existing research software report ChP volume. FreeSurfer, one of the most popular tools for segmenting cortical and subcortical regions in the brain, reports ChP estimates, but recent work has revealed severe inconsistency and inaccuracy in its volume estimates^15–18^. **Figure 2** shows an example of under-segmentation of ChP by FreeSurfer. There are published neuropsychiatry imaging studies which found ChP changes in disorders based on FreeSurfer output^9,13,19^, and the reported association between ChP volume and disorders are called into question.

The inconsistencies in FreeSurfer-derived ChP labels also propagate to some extent to newer methods trained on these labels, such as a newer, brain image processing software called FastSurfer^20,21^. Several groups have therefore bypassed FreeSurfer-derived material and developed new segmentation tools for the lateral ventricle ChP using machine learning techniques, particularly deep-learning models^18,22,23^ and Gaussian mixture models^24^. The datasets used to train these models tend to be modest (N=50, 55, 128 and 72, respectively), from a single scanner, and they targeted individual pathology of interest without extensive coverage on subject age. These characteristics limit the applicability of the published tools on other populations that differ from the training domain. To create a deep learning-based tool which can accurately and reliably segment human lateral ventricle ChP regardless of the image source, the training datasets need to cover a wide range of subject and scanner characteristics.

Here we introduce DataPlex, a large image collection consists of a comprehensive (N=700) bank of T1-weighted brain scans where lateral ventricle ChP had been manually labeled and quality controlled by neuroanatomy experts. DataPlex comprises MRI scans spanning four age groups (children, young adults, adults, and elderly), three scanner manufacturers (Siemens, GE, and Philips), three image resolutions (0.7, 0.8, and 1.0mm isotropic), and multiple neuropsychiatric disorders. We maintained balanced demographic distributions within each scanner-age strata. This carefully balanced dataset was built with the intended purpose of training deep learning algorithms for automatic segmentation. The wide variety of subject characteristics is expected to boost the generalizability of deep learning-based tools trained with DataPlex. When automatic segmentation tools are no longer narrowly specific to a certain age group, scanner or clinical population, quantifying ChP volume in a large (1000+) number of subjects will become highly feasible. Further, when automatic segmentation tools eliminate speed and accuracy as limiting factors for ChP morphology research, opportunities will open for multiple lines of inquiry which are not yet feasible, including the creation of normative curves for healthy ChP development across the lifespan which will require population-level segmented data.

To establish the rigor and consistency of the dataset, below we present a detailed description of DataPlex construction.

## Methods

DataPlex is a collection of 700 T1-weighted MRI images accompanied by manually annotated segmentation mask of the lateral ventricle ChP. All segmentations were created by trained human raters and underwent rigorous multi-stage quality control by expert human raters. DataPlex was designed to address two key limitations of existing ChP segmentation tools: 1) the small size of their training datasets, and 2) limited coverage regarding demographics, imaging protocols, scanner hardware, and clinical populations. These limitations can reduce model robustness and lead to poor generalizability to out-of-distribution settings. To maximize the development of robust, accurate, and generalizable segmentation models, the 700 T1-weighted scans included in DataPlex were carefully curated. Dataplex contains a balanced spread between scans acquired by Siemens, General Electric (GE), and Philips MRI scanners; the subjects are well-represented between ages 6 to 90 years, balanced between male and female, and include healthy controls as well as a variety of neuropsychiatric disorders. Clinical phenotypes represented in DataPlex include mood disorders, psychosis spectrum disorders, Alzheimer’s disease, epilepsy, diabetes, obsessive-compulsive disorders, anxiety, and autism spectrum disorders. To accommodate the field’s shift toward high-resolution neuroimaging, we selected scans from more recent studies at higher resolutions (0.7, 0.8mm and 1mm isotropic) instead of the images at a traditional 1.2mm or 1.5mm level resolution.

In DataPlex, 351 of the 700 T1-weighted scans (approximately 50%) have a corresponding T2-weighted scan acquired during the same imaging session. These T2-weighted images provide complementary anatomical information that support the validation of ChP segmentations on T1-weighted scans, particularly by helping distinguish ChP tissue from nearby periventricular vascular structures. However, because T2-weighted sequences are not routinely available in many clinical and research datasets, requiring paired T1- and T2-weighted scans for all subjects would have substantially reduced the diversity and real-world applicability of DataPlex. We therefore adopted a hybrid design in which multimodal information was incorporated whenever available. The T2-weighted scans served as an anatomical reference for the expert raters, helping them identify complex periventricular vascular patterns, including the striatal-thalamic and lateral veins, during review of the T1-based segmentations. This process supported consistent recognition of subtle anatomical boundaries and allowed the raters to apply the same criteria to scans for which only T1-weighted images were available. The resulting design preserves the utility of DataPlex for the development of T1-weighted segmentation models while ensuring that the annotation protocol is grounded in multimodal anatomical validation, wherever possible. Detailed protocols for annotating and quality-controlling ChP segmentations on T1 scans, with and without accompanying T2 images, are provided in the following subsections.

More than 50 publicly available studies with MRI components were considered, and the initial image pool totaled to more than 15,000 scans. Twenty-nine contributed to the final composition of DataPlex due to appropriate subject age groups, scanners used, and clinical profiles. The 29 source studies are listed in **Table 1**, and the demographic information of the 700 subjects are summarized in **Table 2**. DataPlex is available for download on National Institute of Mental Health’s (NIMH) Data Archive (NDA; https://nda.nih.gov; please see *Data Availability and Usage Statement* below). Public sharing of DataPlex will provide a standard benchmark for training deep learning-based ChP segmentation tools for future research.

**Table 1.**
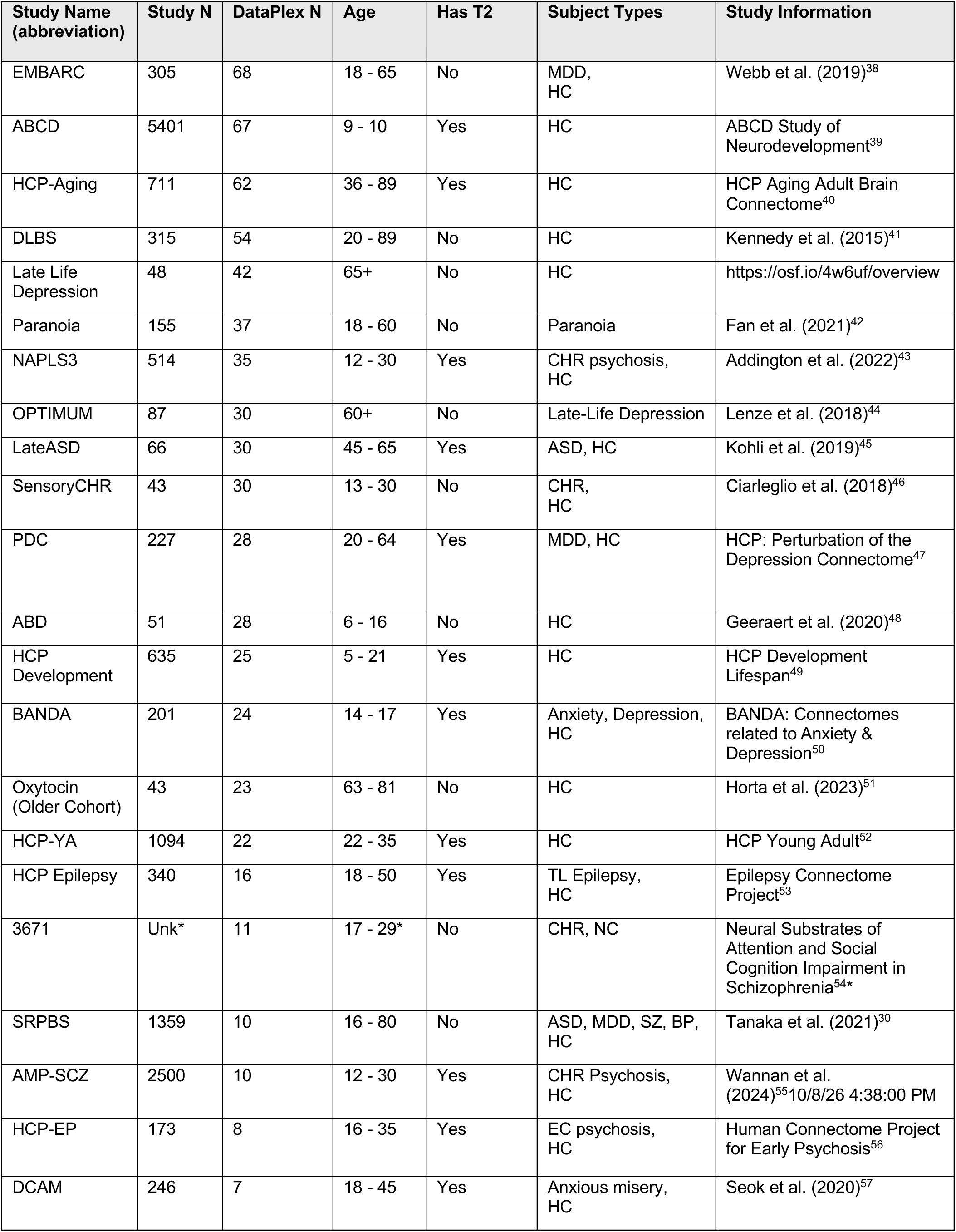
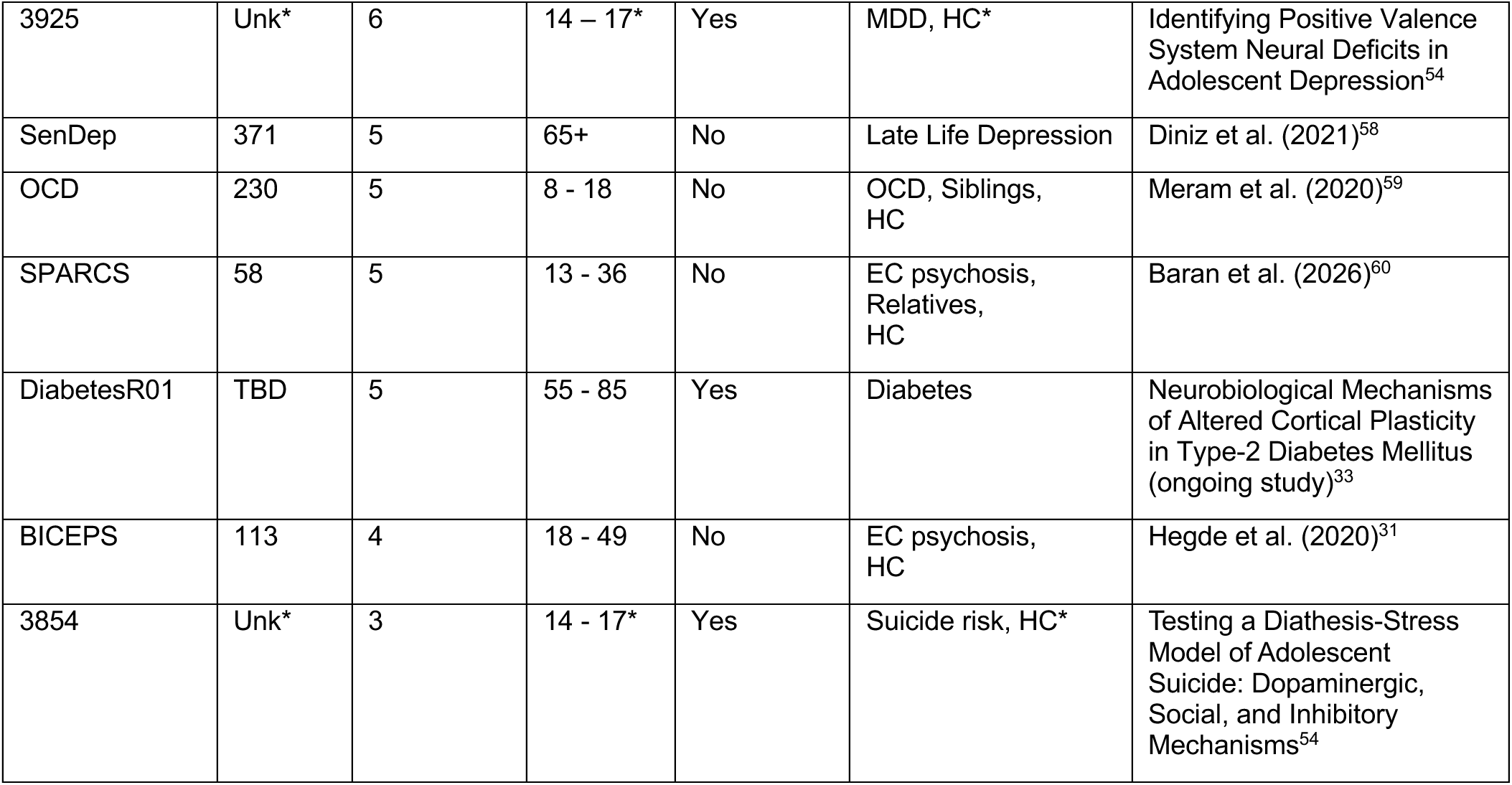
A comprehensive list of studies which contributed to the final DataPlex collection. The number of scans included from each study and subject characteristics are also summarized. ASD: autism spectrum disorder; BP: bipolar disorder; CHR: clinical high-risk for psychosis; EC: early-course psychosis; HC: healthy controls; HCP: Human Connectome Project; MDD: major depressive disorder; OCD: obsessive-compulsive disorder; SZ: schizophrenia. *Three studies (NDA Collection #3671, 3854, and 3925) became unavailable on NDA after we downloaded the images and subject-level demographics. Advanced metadata (clinical diagnoses and race) for subjects in these studies became inaccessible for our team when this table was constructed. The information labeled with asterisks were deduced based on the included subset of scans and the original grant submission for these subjects.

| Study Name (abbreviation) | Study N | DataPlex N | Age | Has T2 | Subject Types | Study Information |
| --- | --- | --- | --- | --- | --- | --- |
| EMBARC | 305 | 68 | 18 - 65 | No | MDD, HC | Webb et al. (2019)<sup>38</sup> |
| ABCD | 5401 | 67 | 9 - 10 | Yes | HC | ABCD Study of Neurodevelopment<sup>39</sup> |
| HCP-Aging | 711 | 62 | 36 - 89 | Yes | HC | HCP Aging Adult Brain Connectome<sup>40</sup> |
| DLBS | 315 | 54 | 20 - 89 | No | HC | Kennedy et al. (2015)<sup>41</sup> |
| Late Life Depression | 48 | 42 | 65+ | No | HC | https://osf.io/4w6uf/overview |
| Paranoia | 155 | 37 | 18 - 60 | No | Paranoia | Fan et al. (2021)<sup>42</sup> |
| NAPLS3 | 514 | 35 | 12 - 30 | Yes | CHR psychosis, HC | Addington et al. (2022)<sup>43</sup> |
| OPTIMUM | 87 | 30 | 60+ | No | Late-Life Depression | Lenze et al. (2018)<sup>44</sup> |
| LateASD | 66 | 30 | 45 - 65 | Yes | ASD, HC | Kohli et al. (2019)<sup>45</sup> |
| SensoryCHR | 43 | 30 | 13 - 30 | No | CHR, HC | Ciarleglio et al. (2018)<sup>46</sup> |
| PDC | 227 | 28 | 20 - 64 | Yes | MDD, HC | HCP: Perturbation of the Depression Connectome<sup>47</sup> |
| ABD | 51 | 28 | 6 - 16 | No | HC | Geeraert et al. (2020)<sup>48</sup> |
| HCP Development | 635 | 25 | 5 - 21 | Yes | HC | HCP Development Lifespan<sup>49</sup> |
| BANDA | 201 | 24 | 14 - 17 | Yes | Anxiety, Depression, HC | BANDA: Connectomes related to Anxiety & Depression<sup>50</sup> |
| Oxytocin (Older Cohort) | 43 | 23 | 63 - 81 | No | HC | Horta et al. (2023)<sup>51</sup> |
| HCP-YA | 1094 | 22 | 22 - 35 | Yes | HC | HCP Young Adult<sup>52</sup> |
| HCP Epilepsy | 340 | 16 | 18 - 50 | Yes | TL Epilepsy, HC | Epilepsy Connectome Project<sup>53</sup> |
| 3671 | Unk* | 11 | 17 - 29* | No | CHR, NC | Neural Substrates of Attention and Social Cognition Impairment in Schizophrenia<sup>54*</sup> |
| SRPBS | 1359 | 10 | 16 - 80 | No | ASD, MDD, SZ, BP, HC | Tanaka et al. (2021)<sup>30</sup> |
| AMP-SCZ | 2500 | 10 | 12 - 30 | Yes | CHR Psychosis, HC | Wannan et al. (2024)<sup>55</sup>10/8/26 4:38:00 PM |
| HCP-EP | 173 | 8 | 16 - 35 | Yes | EC psychosis, HC | Human Connectome Project for Early Psychosis<sup>56</sup> |
| DCAM | 246 | 7 | 18 - 45 | Yes | Anxious misery, HC | Seok et al. (2020)<sup>57</sup> |
| 3925 | Unk* | 6 | 14 – 17* | Yes | MDD, HC* | Identifying Positive Valence System Neural Deficits in Adolescent Depression<sup>54</sup> |
| SenDep | 371 | 5 | 65+ | No | Late Life Depression | Diniz et al. (2021)<sup>58</sup> |
| OCD | 230 | 5 | 8 - 18 | No | OCD, Siblings, HC | Meram et al. (2020)<sup>59</sup> |
| SPARCS | 58 | 5 | 13 - 36 | No | EC psychosis, Relatives, HC | Baran et al. (2026)<sup>60</sup> |
| DiabetesR01 | TBD | 5 | 55 - 85 | Yes | Diabetes | Neurobiological Mechanisms of Altered Cortical Plasticity in Type-2 Diabetes Mellitus (ongoing study)<sup>33</sup> |
| BICEPS | 113 | 4 | 18 - 49 | No | EC psychosis, HC | Hegde et al. (2020)<sup>31</sup> |
| 3854 | Unk* | 3 | 14 - 17* | Yes | Suicide risk, HC* | Testing a Diathesis-Stress Model of Adolescent Suicide: Dopaminergic, Social, and Inhibitory Mechanisms<sup>54</sup> |

**Table 2.** Subject characteristics for the 700 structural MRI volumes included in DataPlex, broken down by age group and scanning parameters (scanner manufacturer and resolution). All voxel sizes listed are isotropic. To fairly represent scanner availability in the field, 36% of DataPlex came from Siemens scanners (N=252), 36% came from GE scanners, and 28% came from Philips scanners (N=196). Note that we were only able to obtain open source 0.7mm scans acquired by Siemens scanners in one age group. Each of the 4 age groups consisted of 175 scans, further balanced by sex within each age group X scanning parameter subcategory. Reference Table 1 for detailed information on the source for these studies. High-resolution (0.7mm isotropic) scans were only available for one age group (young adults). No accessible high-resolution datasets were available for the Philips scanners at the time of DataPlex construction. ASD: autism spectrum disorder; BP: bipolar disorder; CHR: clinical high-risk for psychosis; EC: early-course; HC: healthy controls; HCP: Human Connectome Project; MDD: major depressive disorder; OCD: obsessive-compulsive disorder; SZ: schizophrenia; Unk: *Three studies (NDA Collection#3671, 3854, and 3925) became unavailable on NDA after we downloaded the images and subject-level demographics. Advanced metadata (clinical diagnoses and race) for subjects in these studies became inaccessible for our team when this table was constructed.

|  |  | Siemens |  |  | GE |  | Philips |
| --- | --- | --- | --- | --- | --- | --- | --- |
|  |  | 1.0mm | 0.8mm | 0.7mm | 1.0mm | 0.8mm | 1.0mm |
| Children | N | 19 | 44 |  | 19 | 44 | 49 |
|  | Ave. Age | 11.8 | 14.3 |  | 12.5 | 13 | 9.5 |
|  | Sex (F, M) | 9, 10 | 22, 22 |  | 10, 9 | 22,22 | 25, 24 |
|  | Race (AA, AS, CA, MR, OTH) | 4, 0, 11, 2, 2 | 4, 2, 35, 1, 2 |  | 2, 2, 14, 0, 1 | 1, 0, 0, 0, 0, *UNK=43 | 7, 1, 39, 2, 0 |
|  | Phenotype (N) | NC (10)<br>CHR (5)<br>OCD (4) | NC (22)<br>GAD (12)<br>MDD (10) |  | NC (9)<br>CHR (10) | CHR (1)<br>NC (28)<br>*Unk (15) | NC (49) |
|  | Source Studies (N) | ABCD (9)<br>NAPLS3 (4)<br>OCD (5)<br>SPARCS (1) | BANDA (24)<br>HCP-D (20) |  | ABCD (9)<br>NAPLS3 (10) | 3671 (1)<br>AMP-SCZ (8)<br>3854 (3)<br>3925 (4)<br>ABD (28) | ABCD (49) |
| Young Adults | N | 19 | 22 | 22 | 19 | 44 | 49 |
|  | Ave. Age | 24 | 22.7 | 27.8 | 23.4 | 23.9 | 24.2 |
|  | Sex (F, M) | 10, 9 | 11, 11 | 11, 11 | 9, 10 | 22, 22 | 24,25 |
|  | Race (AA, AS, CA, MR, OTH) | 6, 0, 7, 5, 1 | 5, 2, 14, 0, 1 | 4, 1, 17, 0, 0 |  | 12, 8, 14, 6, 0 | 4, 2, 10, 3, 0 |
|  | Phenotype (N) | NC (9)<br>ECP (1)<br>CHR (6)<br>MDD (3) | NC (12)<br>ECP (4)<br>MDD (6) | NC (22) | NC (6)<br>CHR (7)<br>MDD (6) | NC (20)<br>CHR (20)<br>*Unk (4) | NC (22)<br>ASD (1)<br>BP (1)<br>MDD (20)<br>SAD (5) |
|  | Source Studies (N) | BICEPS (1)<br>EMBARC (4)<br>NAPLS3 (10)<br>SPARCS (4) | DCAM (7)<br>HCP-D (5)<br>HCP-EP (8)<br>PDC (2) | HCP-YA (22) | EMBARC (8)<br>NAPLS3 (11) | SenDep (30)<br>3671 (10)<br>AMP-SCZ (2)<br>3925 (2) | EMBARC (24)<br>Oxytocin (1)<br>Paranoia (24) |
| **Adults** | **N** | 19 | 44 |  | 19 | 44 | 49 |
|  | **Ave. Age** | 50.2 | 46.1 |  | 43.8 | 50.9 | 49.9 |
|  | **Sex (F, M)** | 9, 10 | 22, 22 |  | 10, 9 | 22, 22 | 25, 24 |
|  | **Race (AA, AS, CA, MR, OTH)** | 8, 1, 10, 0, 0 | 6, 4, 31, 2, 1 |  | 3, 10, 6, 0, 0 | 5, 4, 33, 0, 2 | 13, 2, 33, 0, 1 |
|  | **Phenotype (N)** | NC (6)<br>ECP (2)<br>MDD (11) | NC (21)<br>MDD (23) |  | NC (7)<br>MDD (12) | NC (20)<br>ASD (10)<br>TLE (14) | NC (31)<br>BP (4)<br>MDD (6)<br>SAD (4)<br>SZ (4) |
|  | **Source Studies (N)** | BICEPS (3)<br>EMBARC (16) | HCP-A (18)<br>PDC (26) |  | OPTIMUM (2)<br>EMBARC (7)<br>SRPBS (10) | LateASD (28)<br>HCP-Epil (16) | DLBS (21)<br>EMBARC (9)<br>Oxytocin (6)<br>Paranoia (13) |
| **Elderly** | **N** | 19 | 44 |  | 19 | 44 | 49 |
|  | **Age** | 71.2 | 74.3 |  | 72.3 | 64.5 | 75 |
|  | **Sex (F, M)** | 10, 9 | 22, 22 |  | 9, 10 | 29, 15 | 24, 25 |
|  | **Race (AA, AS, CA, MR, OTH)** | 3, 1, 15, 0, 0 | 5, 7, 30, 0, 2 |  | 2, 0, 16, 0, 1 | 0, 42, 2, 0, 0 | 1, 0, 48, 0, 0 |
|  | **Phenotype (N)** | NC (3)<br>MDD (16) | NC (44) |  | NC (3)<br>MDD (14)<br>T2DM (2) | NC (38)<br>ASD (1)<br>Diabetes (4) | NC (49) |
|  | **Source Studies (N)** | 3130 (5)<br>OPTIMUM (14) | HCP-A (44) |  | OPTIMUM (14)<br>DiabetesR01 (5) | Late ASD (2)<br>Late Life Depression (40) | DLBS (33)<br>Oxytocin (16) |

### Data Sources

T1-weighted images with slice thicknesses of 1mm or below were downloaded from open-source research databases including the NDA^25^ (12 studies), the Human Connectome Project^26^ (HCP) consortia (7 studies), the Adolescent Brain Cognitive Development^27^ (ABCD) consortium (1 study), OpenNeuro^28^ (4 studies), Open Science Foundation (OSF)^29^ (1 study), and Japan’s Multi-disorder Imaging Dataset called SRPBS^30^ (1 study). All necessary database-and study-specific Data Use Agreements (DUA) were completed prior to data downloads.

Three additional studies, conducted by colleagues at Beth Israel Deaconess Medical Center (BIDMC) were also included. Among the three, BICEPS^31^ and SPARCS^32^ (please see **Table 1** for more information) from Dr. Keshavan’s laboratory contributed to the young adult and adult samples, and an ongoing study, Diabetes R01^33^ from Dr. Fried’s laboratory contributed to the elderly sample. Image transfer from these colleagues to the segmentation team at University of Rochester Medical Center (URMC) was approved by both institutions via formal DUAs.

### Ethics Statement

All source studies obtained informed consent from subjects for de-identified data sharing. To continue subject protection and confidentiality, images and metadata did not contain any personal health information (PHI) when our team downloaded them onto secure servers at BIDMC and later at URMC when the PI, Dr. Lizano, moved institutions. The segmentation team started manual labeling of DataPlex in early 2024 at BIDMC in Boston, Massachusetts, USA, then the effort moved to URMC in Rochester, NY, USA in late 2025 following a laboratory transition. DataPlex creation was approved by the Institutional Review Boards (IRB) at both institutions as a secondary data inquiry.

### DataPlex Characteristics

To allow for future development of deep learning-based automatic tools for image segmentation, our sampling for DataPlex aimed to produce an image dataset with wide and balanced distributions of subject and scanner characteristics. The resulting set of 700 images were sampled into the following strata from the larger pool of 15,000+ images (**Figure 3**):

- 4 levels for age:

◦ Children and adolescents: 6 to 17 years old
◦ Young adults: 18 to 34 years old
◦ Adults: 35 to 64 years old
◦ Elderly: 65 to 90 years old (subjects over 90 were not included to prevent the risk of identification given the smaller population in this age pool)
- 2 levels for sex: male and female
- 5 levels for race: Caucasian, Asian, Black, other, and multi-race
- 3 levels for scanner manufacturer: Siemens, GE, and Philips
- 3 levels for image resolution: voxel sizes of 1.0mm, 0.8mm, and 0.7mm
- 11 levels for psychopathology:

◦ Healthy controls
◦ Bipolar disorder (BP)
◦ Major depressive disorder (MDD)
◦ Schizophrenia (SZ)
◦ Schizoaffective disorder (SAD)
◦ Autism spectrum disorder (ASD)
◦ Early course psychosis (ECP)
◦ Clinical high-risk for psychosis (CHR)
◦ Obsessive-compulsive disorder or behavior (OCD)
◦ Diabetes

**Figure 3.**
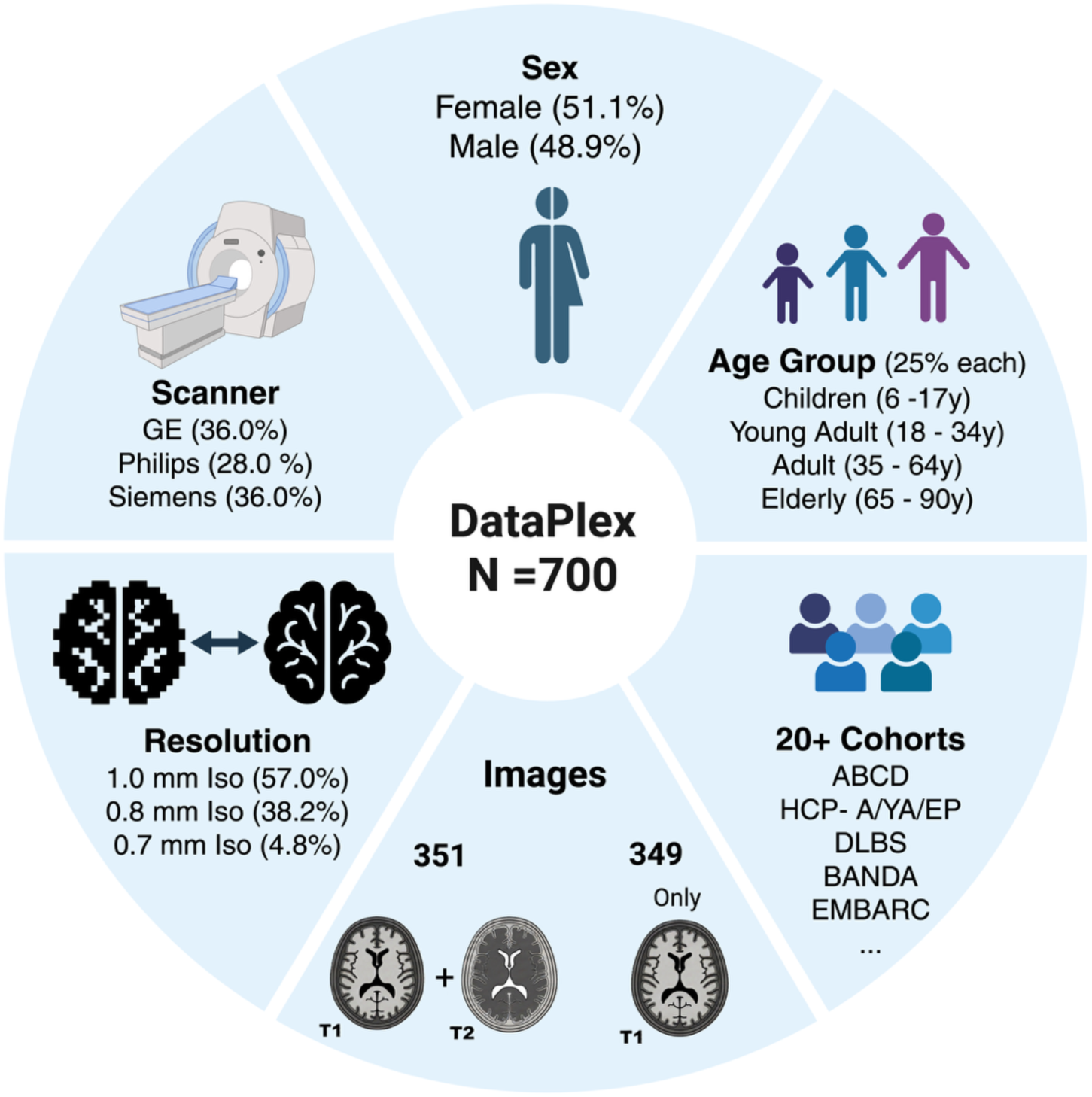
Graphical summary of DataPlex constrction.

We would like to note that infants and newborns were excluded due to the high heterogeneity inherent in this stage of development, and there is currently no gold standard training material for infant lateral ventricle ChP labeling. The final DataPlex sample includes brain images and segmentation masks from subjects between ages 6-90.

### Identifying ChP on T1-weighted Images and Manual Segmentation

Manual segmentation of lateral ventricle ChP was performed using 3D Slicer (version 5.2.2, www.slicer.org). Human raters followed the standardized steps below, also illustrated by step-by-step screenshots in **Supplementary Figure S1**.

1) Load the T1 image in Slicer 5.2.2. This version of 3D Slicer displays MRI images in radiological orientation and allows segmentation masks to match the image’s natural voxel space. Loading a 3D anatomical image populates the three slice views in its graphical user interface (GUI) – coronal, axial, and sagittal views – with centered image slices (**Figure S1.1**).
2) Create two empty segmentation masks, one for the lateral ventricle in the left hemisphere, and one for the lateral ventricle in the right hemisphere (**Figure S1.2**).
3) Adjust brightness and contrast settings for the T1 slice views, focusing on the axial view, to optimize tissue differentiation:

- Ensure the superior portions of the ventricles appear dark. This step leverages the high-water content region to anchor the lower intensity bound, helping suppress low-contrast noise.
- Ensure the internal capsule, which is white-matter dense, is clearly distinguishable from the surrounding striatal nuclei which are grey-matter dense. Maximize the contrast between these two adjacent tissues without overexposing the nearby splenium portion of corpus callosum (**Figure S1.3**). This leverages T1-weighted scans’ inherent property of contrasting lipid-rich versus cell-dense tissue types to 1) ensure the display reflects maximum anatomical fidelity and 2) standardize the visual experience across all raters in a reproducible way.
4) Turn off the paint fill in the segmentation masks (**Figure S1.4**).
5) Select the paint brush; reduce brush size to 1% (**Figure S1.5**).
6) Start segmentation of one hemisphere by anchoring the first labeled voxels on the apical edge of ChP on the axial view (**Figure S1.6**).
7) Slicer 5.2.2 allows voxel co-registration in the interactive GUI - the same voxel can be identified on all 3 views in the GUI simultaneously. Locate the anchored and labeled apical voxels from step 6 in the sagittal view, then use the “paint” function to label the rest of the continuous ChP tissue in the sagittal view (**Figure S1.7**).
8) After labeling all visible ChP voxels, scroll through the labeled area in *sagittal view* in continuous motion and correct missing edges, orphan voxels, and holes using the paint and eraser functions (**Figure S1.8**).
9) Scroll through the labeled area in the *axial view*. Since labeling is complete, although done in a different view, the labeled borders should reflect the visible borders of ChP in this view as well. Use paint and eraser functions to further correct edges, orphan voxels and holes. Use the co-registering feature in 3D Slicer (hold down Shift + cursor movement) to examine voxels of interest and the surrounding anatomy in all 3 axes simultaneously.
10) Finalize the segmentation for the right hemisphere by scrolling through the *coronal view*, again utilizing the co-registering tool. This ensures the labeled ChP boundaries are checked from all 3 angles thoroughly.
11) The anteroinferior boundary of the lateral ChP is structurally defined by its approach to the Foramen of Monro. To prevent spatial encroachment into subsequent segmentations of the third ventricle ChP, the inferior margin of the fornix, clearly identifiable on both coronal and sagittal planes, was established as a strict, unambiguous anatomical boundary. This applied a standardized, reproducible constraint for the extent of our lateral ventricle ChP labels, which ends on the uniform geometric plane intersecting this landmark, isolating the lateral plexus proximal to the narrowest convergence of the interventricular foramen (**Figure. S1.9**).
12) Repeat steps 6-11 to segment and check the ChP in the other hemisphere.
13) Save the two segmentation masks.

This protocol has built-in measures to maximize inter/intra-rater reliability and anatomy accuracy by leveraging the inherent strength of T1-weighted scanning sequences. By maximizing the visual contrast between aqueous regions, grey matter, and white matter in the T1 images, this protocol boosts the signal-to-noise ratio to aid the boundary recognition of the ChP. By standardizing steps 1-3, this protocol maintains a consistent constraint on slice rendering across raters and across scans, supporting reliable segmentation across individuals and time. Steps 6-11 impose a fixed labeling sequence, enforcing a common boundary definition for the lateral ventricle ChP and incorporating a built-in quality check to minimize mislabeling.

After trained raters completed their segmentations, expert raters performed a secondary quality check. Expert raters (LH, NR, and VZ) all had at least two years of experience and maintained high temporal internal consistency. They also maintained high intra-rater consistency with each other. Detailed training protocol for raters is provided in the *Technical Validation* section below, and the training results are shown in **Tables 3, 4, and 5**. During quality control, expert raters followed the same step 3 (**Figure S1.3**) to set up their view of the anatomical image independent of the first rater’s settings. Then the expert raters evaluated the similarity of the ChP boundaries labeled by the first rater against their own view. Edits were made to the segmentation masks completed by the trained rater if necessary. Feedback was provided if the trained raters were found to deviate from the standard visual settings and labeling protocol. This allows a continuous maintenance of between-rater consistency throughout the project.

**Table 3.** Inter-rater reliability among trained raters during DataPlex. The lower triangle (blue) displays Dice coefficients, and the upper triangle (orange) displays the 95% Hausdorf distances in millimeters. A 95% Hausdorf distance of 1.3mm indicates that 95% of the boundary voxels in the two segmentations being compared are between 1 and 2 voxels apart, depending on voxel size. To qualify as a trained rater, the student’s ChP labeling must achieve Dice coefficients >= 0.8 <u>with the expert raters’</u> segmentations (consensus between LH and PL) on three or more training images consecutively. With this entry standard, Table 3 demonstrates coherence in boundary criteria across raters in addition to being independently consistent with expert raters. Reported here are metrics in their last training image, for some it was a 1mm isotropic image, and for others it was a 1.1mm isotropic image. Each trained rater contributed at least 40 segmentations in DataPlex.

|  | AA | BS | DD | DY | EO | MH | NT | PP | VL | WM |
| --- | --- | --- | --- | --- | --- | --- | --- | --- | --- | --- |
| AA |  | 1.34 | 1.27 | 1.12 | 1.20 | 1.13 | 1.00 | 1.12 | 1.05 | 1.05 |
| BS | 0.81 |  | 1.32 | 1.12 | 1.13 | 1.33 | 1.00 | 1.12 | 1.12 | 1.05 |
| DD | 0.75 | 0.78 |  | 1.27 | 1.12 | 1.12 | 1.00 | 1.27 | 1.00 | 1.27 |
| DY | 0.85 | 0.85 | 0.81 |  | 1.05 | 1.13 | 1.40 | 1.12 | 1.12 | 1.05 |
| EO | 0.79 | 0.83 | 0.83 | 0.83 |  | 1.20 | 1.00 | 1.12 | 1.13 | 1.05 |
| MH | 0.82 | 0.76 | 0.76 | 0.78 | 0.75 |  | 1.00 | 1.12 | 1.35 | 1.33 |
| NT | 0.86 | 0.82 | 0.86 | 0.76 | 0.83 | 0.79 |  | 1.33 | 1.00 | 1.27 |
| PP | 0.83 | 0.83 | 0.80 | 0.86 | 0.80 | 0.81 | 0.76 |  | 1.12 | 1.13 |
| VL | 0.82 | 0.77 | 0.76 | 0.79 | 0.78 | 0.76 | 0.80 | 0.80 |  | 1.13 |
| WM | 0.86 | 0.87 | 0.80 | 0.86 | 0.82 | 0.75 | 0.76 | 0.85 | 0.82 |  |

**Table 4.** Inter- and intra-rater reliabilities (IRR) of expert raters’ manual segmentations using T1 images only, measured in Dice coefficients (blue) and 95% Hausdorff distances in mm (orange). Shaded cells indicate longitudinal IRR within each expert rater. Intra-rater reliability (central diagonal) was obtained by each expert rater independently segmenting the same MRI volume at two different time points, at least 3 months apart. Inter-rater reliability was measured via pairs of expert raters segmenting the same MRI volume within a week.

|  | Expert Rater 1 (LH) |  | Expert Rater 2 (NR) |  | Expert Rater 3 (VZ) |  |
| --- | --- | --- | --- | --- | --- | --- |
| Expert Rater 1 | 0.90 | 1.05 | 0.97 |  | 0.8 |  |
| Expert Rater 2 | 0.84 |  | 0.92 | 1.00 | 1.00 |  |
| Expert Rater 3 | 0.91 |  | 0.87 |  | 0.89 | 0.99 |

**Table 5.** Inter- and intra-rater reliabilities (IRR) of expert raters’ QC edit results using T1 and T2 scans, measured in Dice coefficients (blue) and average Hausdorff distances in mm (orange). Shaded cells indicate longitudinal IRR within each expert rater. Intra-rater reliability (central diagonal) was obtained by each rater independently making QC edits on the same manual segmentation at two different time points, at least 3 months apart. Inter-rater reliability was measured via pairs of expert raters making edits on the same segmentation. Average Hausdorff distances are displayed in this table because the 95% Hausdorff distances were mostly 0 and not as informative.

|  | Expert Rater 1 (LH) |  | Expert Rater 2 (NR) |  | Expert Rater 3 (VZ) |  |
| --- | --- | --- | --- | --- | --- | --- |
| Expert Rater 1 | 0.96 | 0.08 | 0.09 |  | 0.07 |  |
| Expert Rater 2 | 0.96 |  | 0.97 | 0.06 | 0.07 |  |
| Expert Rater 3 | 0.98 |  | 0.96 |  | 0.84 | 0.30 |

### Additional Segmentation Protocol for Images Accompanied by a T2

Out of the 700 T1-weighted brain scans in DataPlex, T2-weighted images acquired during the same scanning session are available in 351 subjects. T2-weighted images are valuable for lateral ventricle ChP segmentation for their ability to highlight vasculature. T1-weighted scanning sequences are designed to distinguish tissues with differing levels of lipid content, where blood vessels often appear with similar intensity as ChP. This created some level of ambiguity in ChP boundaries when only T1 images were available. Some transverse veins unrelated to the immediate ChP anatomical complex, such as the lateral veins and striatal-thalamic veins, are in the vicinity of lateral ventricle ChP and are therefore difficult to parse on T1 images without extensive experience (**Supplementary Figure S2**, top row). T2-weighted images can capture fast-flowing liquid; the speed of fluid movement in blood vessels far surpasses movement in the ventricles, and the flow creates a signal void effect resulting in dark vascular patterns in T2 images. The stark signal void is clearly visible and easily differentiates the vasculature from the nearby ChP, providing unique anatomical insight aiding ChP segmentation (An example is shown in **Figure S2**, bottom row). For the 351 entries in DataPlex where T1 and T2 images were both available, the two images were spatially co-registered using `bbregister`, which is part of the FreeSurfer 7.1.0 suite^34^.

Definitions of trained raters, expert raters, and quality control are described in the Technical Validation section below. Briefly, after trained raters completed manual segmentations based on the T1 image, expert raters used the co-registered T1 and T2 images to perform quality control checks and edits. They incorporated the additional anatomical boundaries informed by T2 images and removed voxel labels which mistakenly captured transverse, non-ChP supplying vasculature.

## Technical Validation

The DataPlex team implemented a multi-layer quality control pipeline to ensure the accuracy and consistency of the 700 manual segmentations. The workflow involved two rater groups, trained raters and expert raters, and two stages of mask generation: an initial pass by trained raters followed by a quality control (QC) pass by expert raters. Research staff were required to complete standardized training and pass a qualification assessment to serve as trained raters. Expert raters were senior team members with at least two years of experience who had completed more than 100 segmentations in the present or previous projects, while maintaining high intra- and inter-rater consistency. The initial training protocol and longitudinal quality validation procedures are described below.

All rater candidates completed segmentations on a set of training images independent of DataPlex, which spanned 0.8, 1.0, 1.1 and 1.2mm resolutions and each selected to present distinct challenges in ChP segmentation (e.g., motion artifacts, atypical ventricular anatomy, narrow ventricles etc.). Reference (“gold standard”) segmentations were generated by expert raters (NR and LH; 4 and 2 years of experience, respectively) and approved by the principal investigator (PL; 10 years of experience). We used Dice Coefficient to measure the level of spatial overlap between trainee vs. expert segmentations (and later for longitudinal intra-rater consistency as well), and the mathematical formula is:

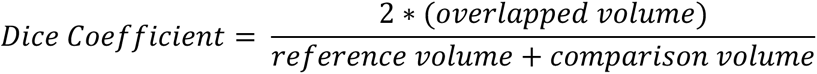

The Dice formula results in a unit-less ratio between 0 and 1 which can quickly summarize the similarity of *placement* between two segmentations. The higher the coefficient, the higher proportion of the two segmentations overlap with each other.

To quantify the variations in the *strictness* of voxel inclusion across raters, we used the 95% Hausdorff Distance as computed by the SlicerRT Segment Comparison module^35^, which reports the mean of the 95^th^-percentile nearest-neighbor distances between the surfaces of reference and comparison volumes. Hausdorff Distances carry the same one-dimensional unit as the measurement of voxel sizes, in this case mm. Inter-rater 95% Hausdorff distances considered “good” in the ChP MRI image segmentation literature range from 1.69 to 2.22mm^22,36^. ChP segmentations by FreeSurfer shared a 95% Hausdorff distance with manual segmentation of over 10mm^22^.

The training process focused on helping raters learn the correct anatomical landmarks for placement. During training, the expert-labeled masks were withheld from the candidates to ensure independent protocol internalization. Aiming for our human raters to outperform the published automatic segmentation tools, which reported the tools sharing agreement with manual segmentation at Dice = 0.65-0.8^18,22,24,36^, DataPlex trainees must score a Dice Coefficient higher than 0.8 for three consecutive training images to qualify as a trained rater. All trained raters met this qualification threshold using the same set of training images before gaining access to DataPlex.

To ensure labeling efficiency and accountability, each trained rater was assigned a unique set of 40 scans from DataPlex. Additional sets were only assigned by expert raters after verifying the quality and completion of the first set. Each trained rater labeled at least one set of images (min = 40, max = 132 images) over the average duration of 3 to 4 months. Continued inter-rater reliability was assessed through 5 additional practice scans scattered across their assignments, at least 6 scans apart. These continued inter-rater reliability (measured in both Dice Coefficients and Hausdorff Distances) among the trained raters are summarized in **Table 3**.

The quality control stage determined the final accuracy of the segmentation masks, so the cross-sectional inter-rater reliability and longitudinal intra-rater reliability among expert raters were even more strictly maintained than among trained raters. Reliability among segmentations performed by expert raters is summarized in **Table 4**. In addition to quantitative measurements of IRRs, **Figure 4** provides a visual example of realistic agreement for lateral ChP segmentation among the DataPlex expert raters.

**Figure 4.**
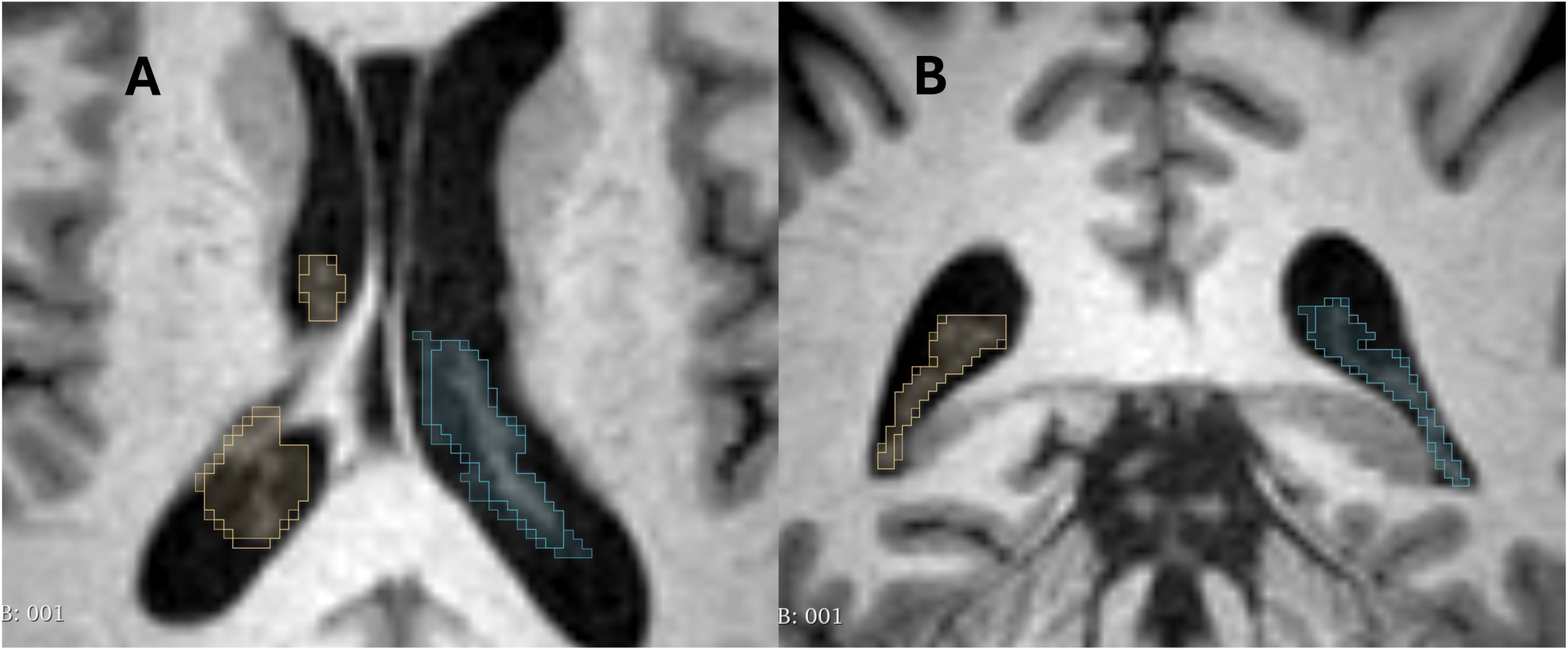
Intra-rater and inter-rater reliabilities are maintained above Dice coefficient > 0.8. This figure gives a representative demonstration of Dice = 0.85, from one expert rater performing segmentation on the same T1 volume twice, 6 months apart. The first and second attempts were overlayed together, with **A** showing similar ChP labeling on the axial view, and **B** showing the two attempts overlayed in the coronal view.

Consistency was also maintained in our tool selection and image preparations. 3D Slicer is an open-source software with an active user base, and the development team updates its core functions frequently, including changing the default orientation for image display between versions. To prevent inconsistencies in the co-registration between the T1 image space, segmentation masks, and anatomical fiducials, all members of the DataPlex team maintained the identical software version (5.2.2) - including all trained raters and all expert raters -throughout the project. Image preparation steps of all 700 scans in DataPlex were identical: they all started with the selected T1 scans in standard NIfTI file format. To be readily displayed and labeled in 3D Slicer, these NifTI images underwent format conversion to MGZ via the FreeSurfer command `mri_convert`. No additional volume processing such as image normalization was performed before manual labeling in 3D Slicer, and each scan retained their original resolution: 0.7, 0.8, or 1.0mm isotropic. In one study, original images were acquired at 0.8mm isotropic by a GE scanner, however the T1-weighted protocol was zero-filled interpolated in-plane so that the resulting resolution was 0.5mm x 0.5mm x 0.8mm. In these cases, `mri_convert` was used to re-slice the image to 0.8mm isotropic before being labeled.

Segmentations completed in 3D Slicer were first saved in its native *seg.nrrd* multi-layer binary format, then 3D Slicer was used to convert the binary masks from *seg.nrrd* to *mgz*. After final QC was completed, *mgz* copies of the segmentation masks were made for storage and repository submissions.

## Data Availability Statement

The data bundle can be downloaded from the study’s collection page on NDA at https://nda.nih.gov/edit_collection.html?id=5175. The bundle includes 700 segmentation masks and a comma-delimited metadata sheet. As the 700 included MRI images were sourced from 29 studies, each adhering to different data use agreement terms, we extensively document the path names, source links, unique GUIDs of each included subject, and the study-specific subject IDs for each scan in the metadata sheet, so future users of DataPlex can independently identify and obtain all the source images.

## Data Records

After unzipping, the folder will contain the following files:

- 700 manually labeled, binary segmentation masks in *mgz* format, matching the image space of the source T1-weighted scans. For a portion of the scans, the manual labeling also referenced co-registered T2-weighted scans when those were available. For details about this subset, please see the “*Additional Segmentation Protocol for Images Accompanied by a T2”* subsection above.
- Metadata sheet compatible with NDA’s image03.csv data structure, which contains information about the T1-weighted scans. Columns include: original study name, NDA global unique identifier (GUID) or pseudo-GUID equivalent, subject age, sex, race, scanner manufacturer, resolution, and whether the segmentation also referenced a T2-weighted scan.

## Author Contributions

PL generated the research idea, obtained funding for this project and trained the expert raters. LH refined the labeling protocol from previous work by PL, DB, and YC, who are experienced expert raters from previous work. LH and VZ drafted the manuscript. LH, NR and VZ were the three expert raters who performed quality checks for DataPlex. MK and PF advised on stratified sampling during DataPlex construction and contributed images following data sharing agreements stated in the subject consent, as well as URMC and BIDMC regulations. MR, DK and SE advised on image processing, image set breakdown and provided parameters and constraints for tailoring the DataPlex to meet machine learning tool development needs. DY, DD, EO, MH, VL, AA, WM, BS, PP, and NT were the trained raters. All authors reviewed, edited, and approved the manuscript prior to submission.

## Acknowledgements and Funding Information

This project is supported by NIMH grant MH131586 to Dr. Lizano (PI) and Dr. Reuter (Co-I). We would like to thank Oshin Kushwaha for the assistance in setting up the project infrastructure. Below we acknowledge and provide the funding information for the original studies which generated the MRI images in DataPlex.

## ABCD

Data used in the preparation of this article were obtained from the Adolescent Brain Cognitive Development (ABCD) Study (https://www.nbdc-datahub.org/), held in the NIMH Data Archive (NDA). This is a multisite, longitudinal study designed to recruit more than 10,000 children age 9-10 and follow them over 10 years into early adulthood. The ABCD Study® is supported by the National Institutes of Health and additional federal partners under award numbers: U01DA041048, U01DA050989, U01DA051016, U01DA041022, U01DA051018, U01DA051037, U01DA050987, U01DA041174, U01DA041106, U01DA041117, U01DA041028, U01DA041134, U01DA050988, U01DA051039, U01DA041156, U01DA041025, U01DA041120, U01DA051038, U01DA041148, U01DA041093, U01DA041089, U24DA041123, U24DA041147. A full list of supporters is available at https://abcdstudy.org/federal-partners.html. A listing of participating sites and a complete listing of the study investigators can be found at https://abcdstudy.org/consortium_members/. ABCD consortium investigators designed and implemented the study and/or provided data but did not necessarily participate in the analysis or writing of this report. This manuscript reflects the views of the authors and may not reflect the opinions or views of the NIH or ABCD consortium investigators. The ABCD data repository grows and changes over time. The ABCD data used in this report came from DOI: 10.15154/z563-zd24.

## AMP-SCZ

Data used in the preparation of this article were obtained from the AMPSCZ study (https://www.ampscz.org), held in the NIMH Data Archive (NDA). This is a multisite, longitudinal study designed to recruit more than 2,500 subjects ages 12 to 30 and follow them over the course of 24 months. The AMP SCZ research program is supported by contributions from the AMP SCZ public and private partners, including the National Institute of Mental Health (NIMH) under award numbers: U24MH124629, U01MH124631, U01MH124639, the Wellcome Trust (220664/Z/20/Z and 220664/A/20/Z), and the NHMRC CRE PRE-EMPT (1198304). A full list of supporters is available at https://www.ampscz.org/about/partners/, and a complete listing of the study investigators is at https://www.ampscz.org/about/research-networks/. The AMP SCZ is the largest international collaboration to date, consisting of 42 sites from 13 participating countries. A complete list of participating sites can be found at https://www.ampscz.org/about/map/. AMP SCZ consortium investigators designed and implemented the study and/or provided data but did not necessarily participate in the analysis or writing of this report. This manuscript reflects the views of the authors and may not reflect the opinions or views of the NIH or AMP SCZ consortium investigators.

## HCP-YA

Data were provided [in part] by the Human Connectome Project, WU-Minn Consortium (Principal Investigators: David Van Essen and Kamil Ugurbil; 1U54MH091657) funded by the 16 NIH Institutes and Centers that support the NIH Blueprint for Neuroscience Research; and by the McDonnell Center for Systems Neuroscience at Washington University.

## HCP-Aging

Research reported in this publication was supported by the National Institute on Aging of the National Institutes of Health under Award Number U01AG052564. The content is solely the responsibility of the authors and does not necessarily represent the official views of the National Institutes of Health. The HCP-Aging 2.0 Release data used in this report came from DOI: 10.15154/1520707.

## HCP-Development

Research reported in this publication was supported by the National Institute of Mental Health of the National Institutes of Health under Award Number U01MH109589 and by funds provided by the McDonnell Center for Systems Neuroscience at Washington University in St. Louis. The HCP-Development 2.0 Release data used in this report came from DOI: 10.15154/1520708.

## SRPBS

Data used in the preparation of this work were obtained from the DecNef Project Brain Data Repository (https://bicr-resource.atr.jp/srpbsopen/), collected as part of the Japanese Strategic Research Program for the Promotion of Brain Science (SRPBS) supported by the Japanese Advanced Research and Development Programs for Medical Innovation (AMED).

## Funding Information for Other Included Studies

Oxytocin: UL1-TR000064; R01AG059809, R24 AG039350, R24 AG054355, 3R01AG072658-01A1S, and T32 AG020499; National Science Foundation Cooperative Agreement No. DMR-1157490; U.S. Navy Office of Naval Research Grant N00014-21-1-2201

NAPLS3 (DOI: 10.15154/304r-9v58): U01MH081928, U01MH081988, U01MH081984, U01MH082004, U01MH081902, U01MH082022, U01MH081944, U01MH076989, U01MH081857

BANDA (DOI: 10.15154/6fng-rj25): U01MH108168

OCD (DOI: 10.15154/aps3-0723): R01MH059299-15

Paranoia (DOI: 10.15154/pmb4-r907): R21MH112930

SPARCS: K01MH114012, R01MH107579-04

BICEPS: R01MH092440

EMBARC (DOI: 10.15154/e99n-np66): U01MH092221, U01MH092250

DLBS: R37AG-006265

DCAM (DOI: 10.15154/6neq-wm52): U01MH109991

PDC (DOI: 10.15154/00b0-hw43): U01MH110008

LateASD (DOI: 10.15154/nr0y-k674): R01MH103494

HCP-Epilepsy: U01NS093650

Diabetes-R01: R01AG070077

3671: K23MH108711

3854: R01MH126181

3925: R21MH125044

SenDep (DOI: 10.15154/z3e6-w042): R01MH115953

OPTIMUM: R01MH114966, R01MH114969, R01MH114970, R01MH114980, R01MH114981

ABD: Natural Sciences and Engineering Research Council Discovery Grants (Canada)

Late Life Depression: The First Affiliated Hospital of Nanchang University Internal Funding

---

**Figure S1.**
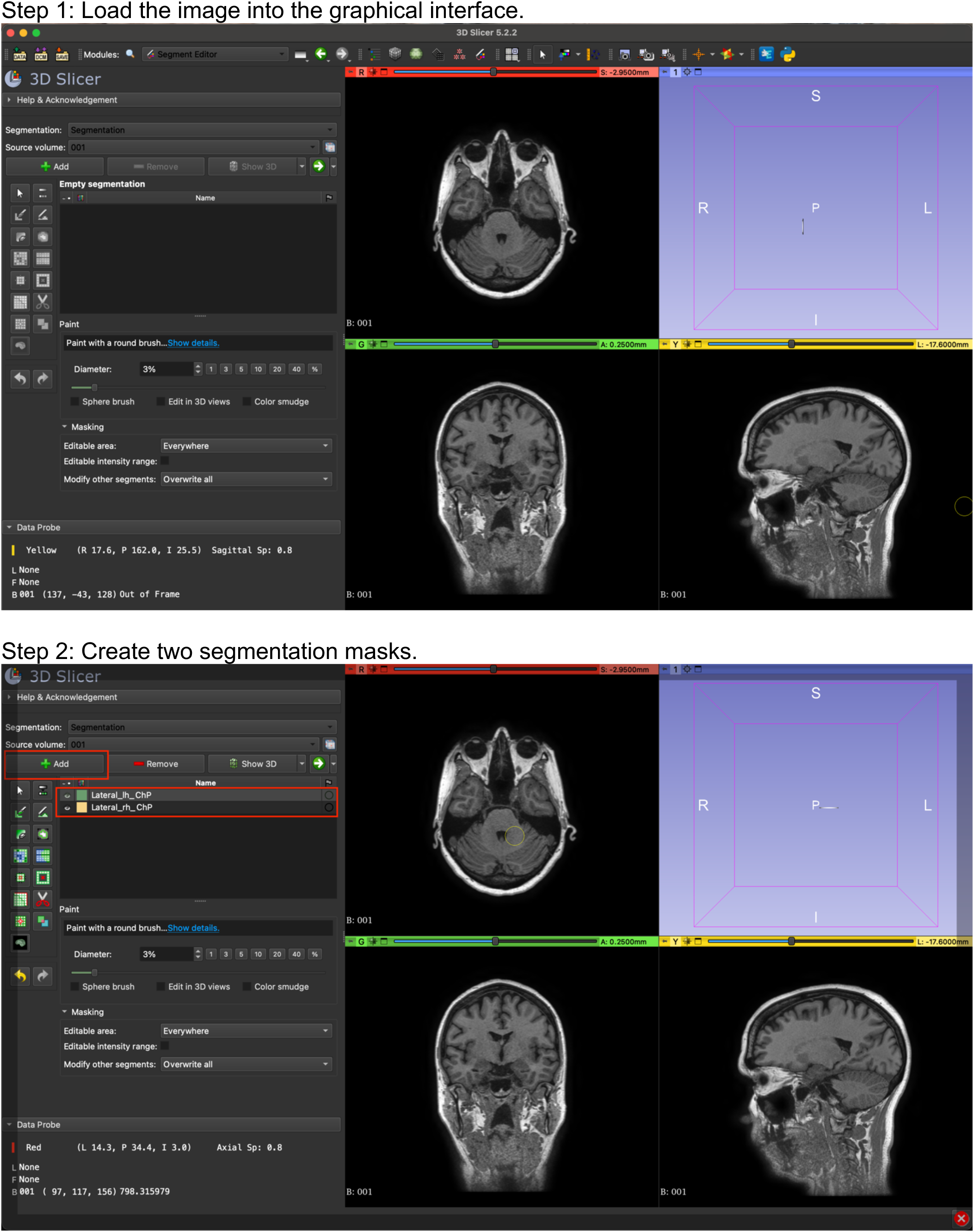
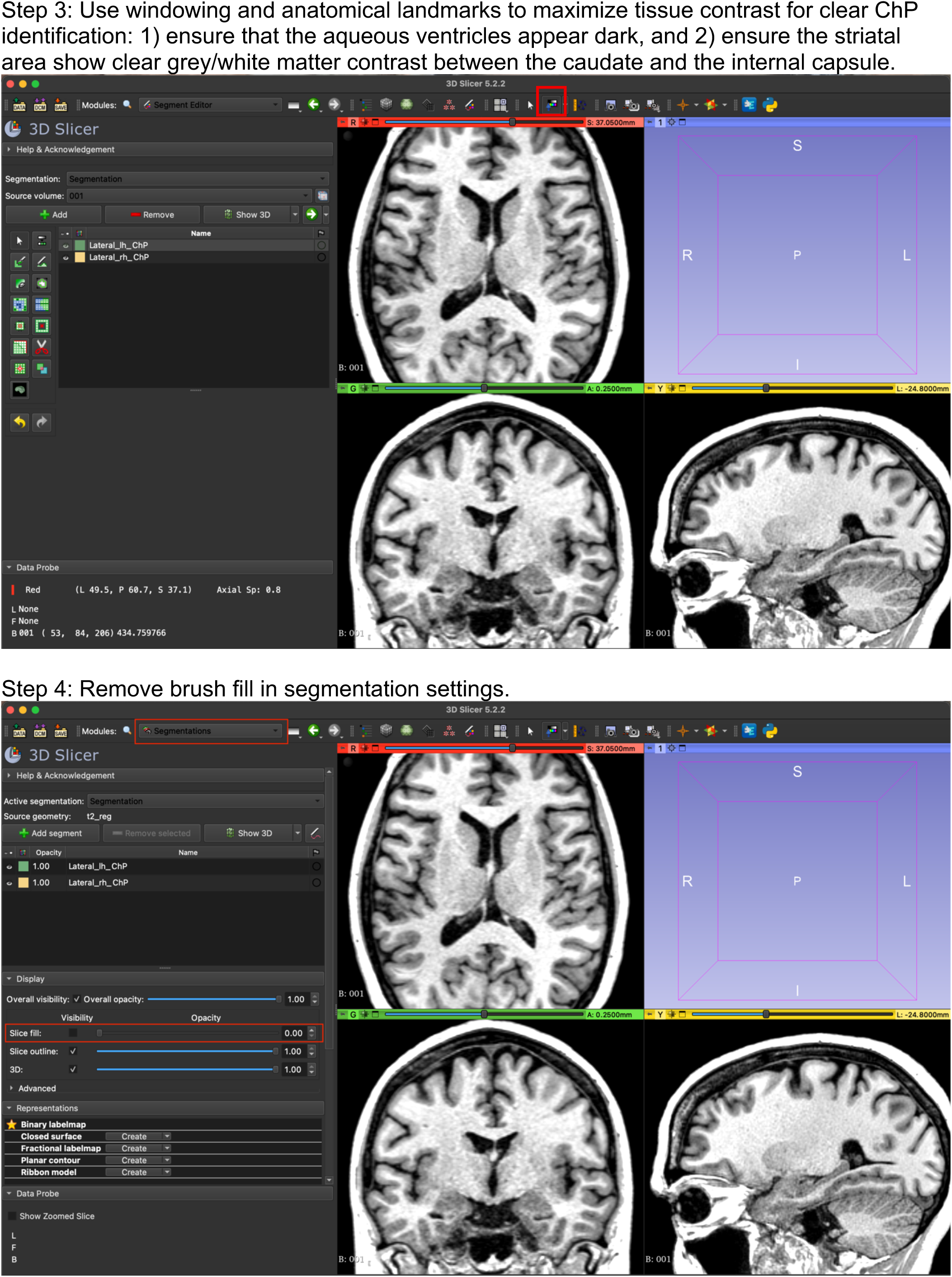
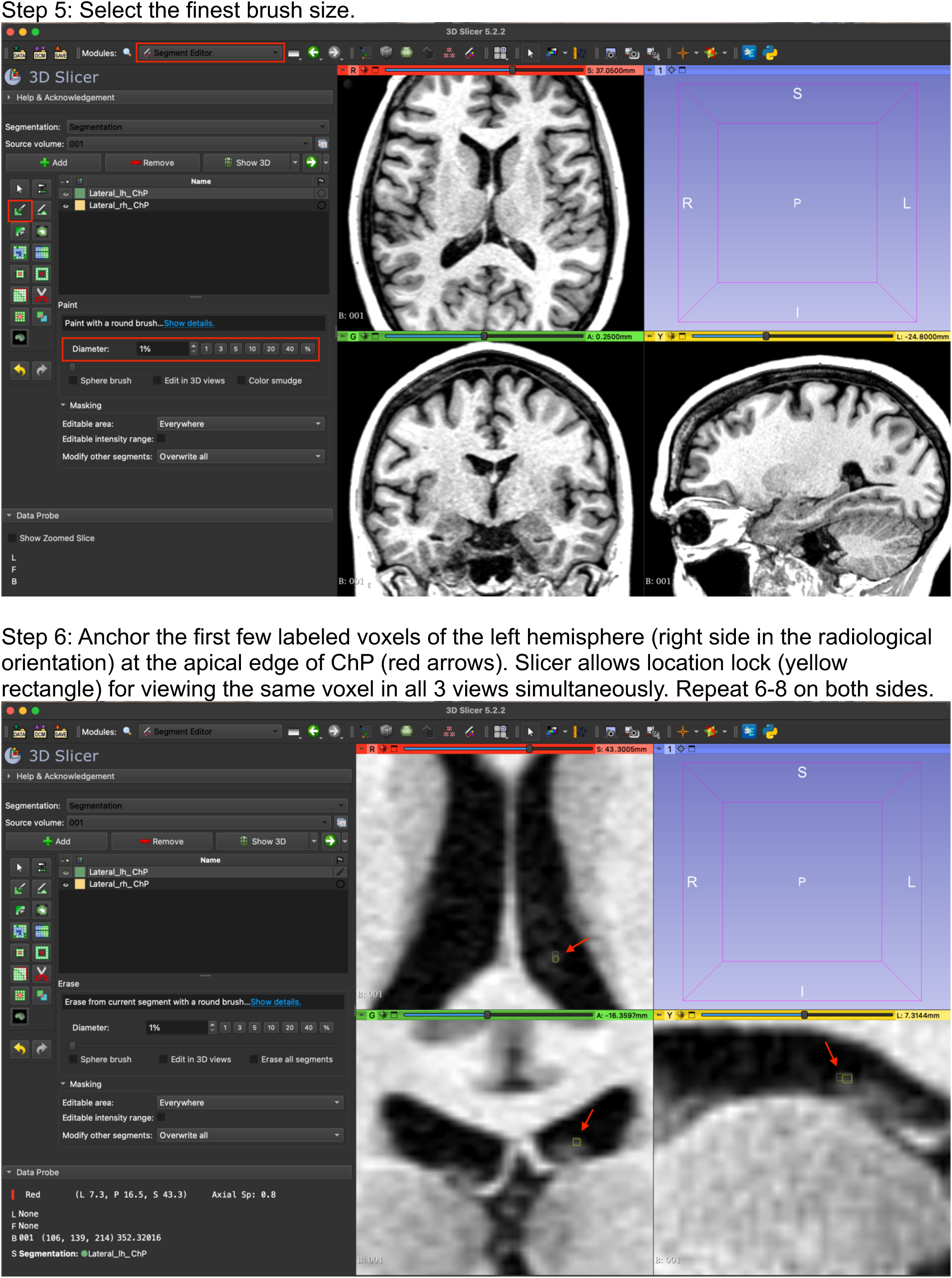
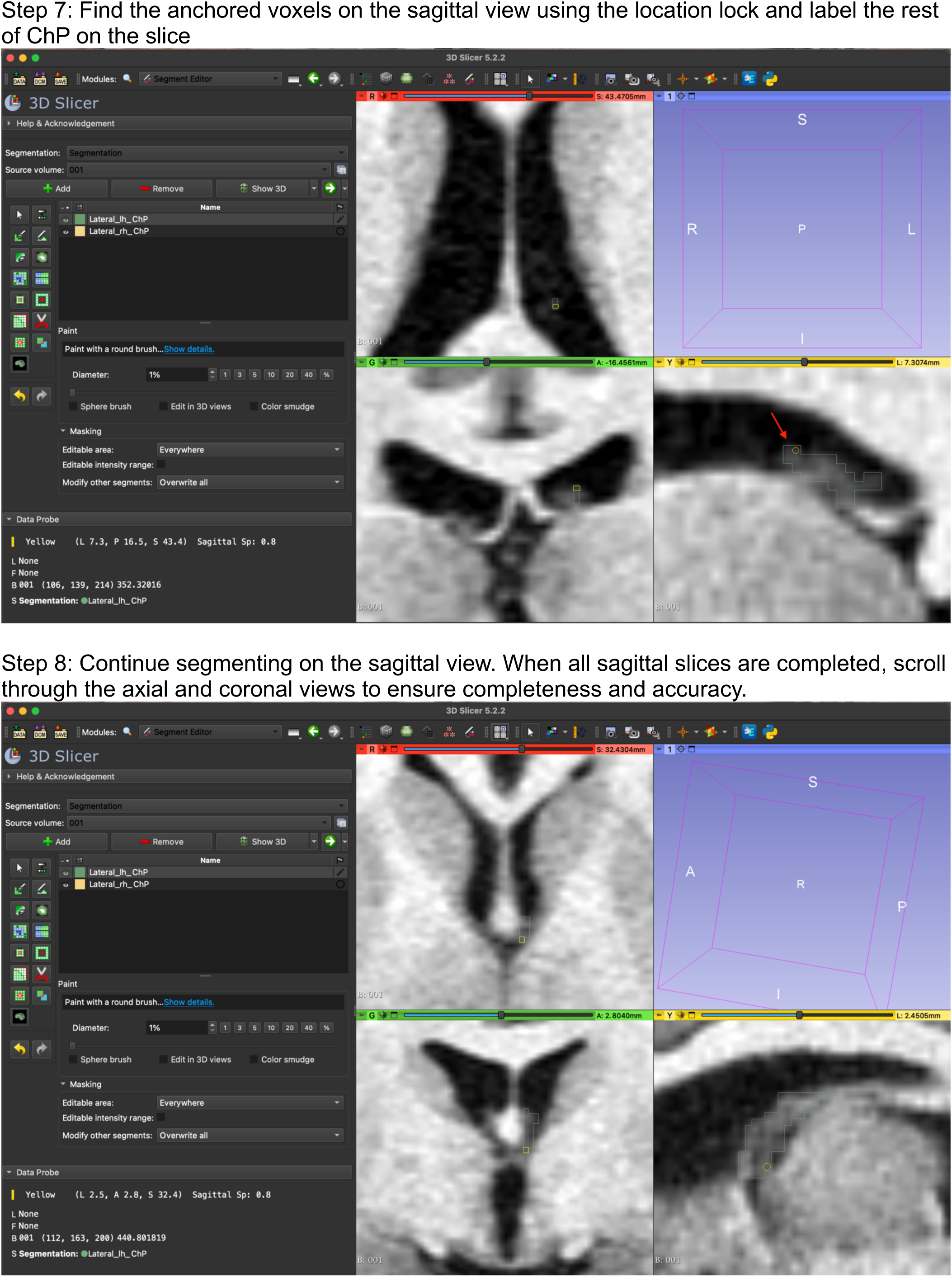
Step-by-step tutorial for manual segmentation of human lateral ventricle ChP using 3D Slicer. This protocol is followed throughout DataPlex construction.

**Figure S2:**
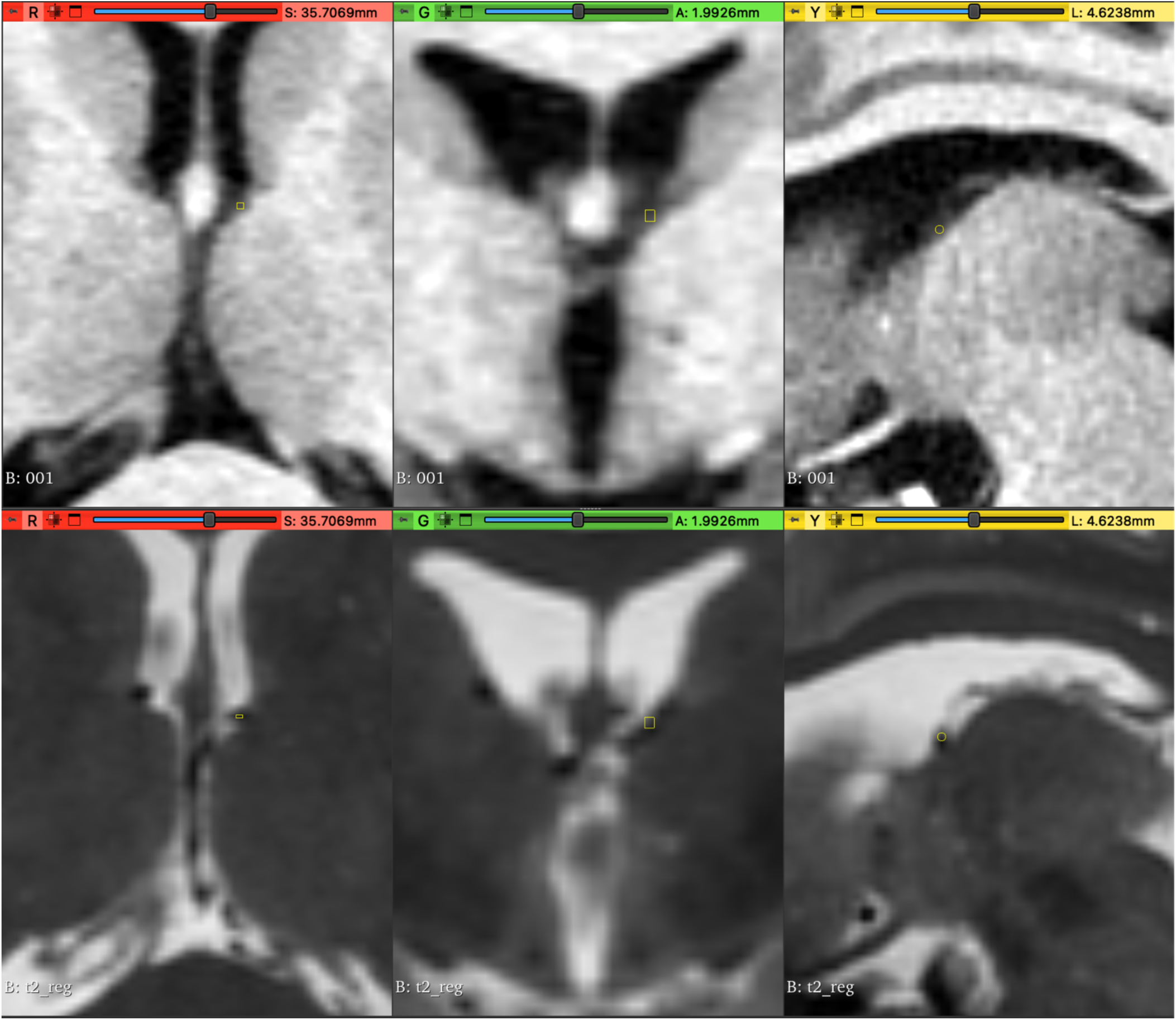
T2-weighted images highlight vasculature and aids manual segmentation of lateral ventricle ChP. This figure shows a direct comparison of T1-weighted and T2-weighted images surrounding the same voxel. The T1 image slices surrounding the same voxel (yellow square, pointing to the striatal-thalamic vein) are displayed in the top row, and the T2 images are on the bottom. Left to right: axial, coronal, and sagittal views.

## References

1. Ding, Z. et al. The glymphatic system: a new perspective on brain diseases. Front. Aging Neurosci. 15, 1179988 (2023).

2. Bitanihirwe, B. K. Y., Lizano, P. & Woo, T. W. Deconstructing the functional neuroanatomy of the choroid plexus: an ontogenetic perspective for studying neurodevelopmental and neuropsychiatric disorders. Mol Psychiatry 27, 3573–3582 (2022).

3. Jessen, N. A., Munk, A. S., Lundgaard, I. & Nedergaard, M. The Glymphatic System: A Beginner’s Guide. Neurochem Res 40, 2583–99 (2015).

4. Plog, B. A. & Nedergaard, M. The glymphatic system in CNS health and disease: past, present and future. Annu. Rev. Pathol. 13, 379–394 (2018).

5. Damkier, H. H., Brown, P. D. & Praetorius, J. Cerebrospinal fluid secretion by the choroid plexus. Physiol Rev 93, 1847–92 (2013).

6. Demeestere, D., Libert, C. & Vandenbroucke, R. E. Therapeutic implications of the choroid plexus–cerebrospinal fluid interface in neuropsychiatric disorders. Brain. Behav. Immun. 50, 1–13 (2015).

7. Fleischer, V. et al. Translational value of choroid plexus imaging for tracking neuroinflammation in mice and humans. Proc. Natl. Acad. Sci. U. S. A. 118, e2025000118 (2021).

8. Jeong, S. H. et al. Choroid Plexus Volume, Amyloid Burden, and Cognition in the Alzheimer’s Disease Continuum. Aging Dis 10.14336/AD.2024.0118 (2024) doi:10.14336/AD.2024.0118.

9. Lizano, P. et al. Association of Choroid Plexus Enlargement With Cognitive, Inflammatory, and Structural Phenotypes Across the Psychosis Spectrum. Am J Psychiatry 176, 564–572 (2019).

10. Althubaity, N. et al. Choroid plexus enlargement is associated with neuroinflammation and reduction of blood brain barrier permeability in depression. NeuroImage Clin. 33, 102926 (2021).

11. Umemura, Y. et al. Choroid plexus enlargement in mild cognitive impairment on MRI: a large cohort study. Eur. Radiol. 34, 5297–5304 (2024).

12. Zhou, Y.-F. et al. Choroid Plexus Enlargement and Allostatic Load in Schizophrenia. Schizophr. Bull. 46, 722–731 (2020).

13. Zhou, G., Hotta, J., Lehtinen, M. K., Forss, N. & Hari, R. Enlargement of choroid plexus in complex regional pain syndrome. Sci. Rep. 5, 14329 (2015).

14. Cao, Y. et al. Brain-derived subgroups of bipolar II depression associate with inflammation and choroid plexus morphology. Psychiatry Clin. Neurosci. 77, 613–621 (2023).

15. Fischl, B. FreeSurfer. NeuroImage 62, 774–781 (2012).

16. Bannai, D., Cao, Y., Keshavan, M., Reuter, M. & Lizano, P. Manual Segmentation of the Human Choroid Plexus Using Brain MRI. J Vis Exp 10.3791/65341 (2023) doi:10.3791/65341.

17. Senay, O. et al. A preliminary choroid plexus volumetric study in individuals with psychosis. Hum. Brain Mapp. 44, 2465–2478 (2023).

18. Visani, V. et al. ASCHOPLEX: A generalizable approach for the automatic segmentation of choroid plexus. Comput. Biol. Med. 182, 109164 (2024).

19. Alisch, J. S. R., Egan, J. M. & Bouhrara, M. Differences in the choroid plexus volume and microstructure are associated with body adiposity. Front. Endocrinol. 13, 984929 (2022).

20. Henschel, L. et al. FastSurfer - A fast and accurate deep learning based neuroimaging pipeline. Neuroimage 219, 117012 (2020).

21. Henschel, L., Kügler, D. & Reuter, M. FastSurferVINN: Building resolution-independence into deep learning segmentation methods—A solution for HighRes brain MRI. NeuroImage 251, 118933 (2022).

22. Eisma, J. J. et al. Deep learning segmentation of the choroid plexus from structural magnetic resonance imaging (MRI): validation and normative ranges across the adult lifespan. Fluids Barriers CNS 21, 21 (2024).

23. Yazdan-Panah, A. et al. Automatic segmentation of the choroid plexuses: Method and validation in controls and patients with multiple sclerosis. NeuroImage Clin. 38, 103368 (2023).

24. Storelli, L. et al. A Fully Automatic Method to Segment Choroid Plexuses in Multiple Sclerosis Using Conventional MRI Sequences. J. Magn. Reson. Imaging 59, 1643–1652 (2024).

25. NIMH Data Archive - NDA. https://nda.nih.gov/.

26. Connectome - Homepage. https://www.humanconnectome.org/.

27. Access & download data. https://docs.abcdstudy.org/latest/usage/access.html.

28. OpenNeuro - MRI Search. https://openneuro.org/search/modality/mri?query=%7B%22modality_selected%22%3A%22mri%22%7D.

29. OSF. https://osf.io/search.

30. Tanaka, S. C. et al. A multi-site, multi-disorder resting-state magnetic resonance image database. Sci Data 8, 227 (2021).

31. Hegde, R. R. et al. Association of white matter microstructure and extracellular free-water with cognitive performance in the early course of schizophrenia. Psychiatry Res. Neuroimaging 305, 111159 (2020).

32. Manoach, D. S. et al. Sleep spindle deficits in antipsychotic-naïve early course schizophrenia and in non-psychotic first-degree relatives. Front. Hum. Neurosci. 8, 762 (2014).

33. Fried, P. RePORT ⟩ RePORTER. https://reporter.nih.gov/search/owY6qUBj_EmEHDLabM9-mg/project-details/11037985.

34. Greve, D. N. & Fischl, B. Accurate and Robust Brain Image Alignment using Boundary-based Registration. NeuroImage 48, 63–72 (2009).

35. Pinter, C., Lasso, A., Wang, A., Jaffray, D. & Fichtinger, G. SlicerRT: Radiation therapy research toolkit for 3D Slicer. Med. Phys. 39, 6332–6338 (2012).

36. Zhao, L., Feng, X., Meyer, C. H. & Alsop, D. C. Choroid Plexus Segmentation Using Optimized 3D U-Net. in 2020 IEEE 17th International Symposium on Biomedical Imaging (ISBI) 381–384 (2020). doi:10.1109/ISBI45749.2020.9098443.

37. Thompson, D., Brissette, C. A. & Watt, J. A. The choroid plexus and its role in the pathogenesis of neurological infections. Fluids Barriers CNS 19, 75 (2022).

38. Webb, C. A. et al. Personalized prediction of antidepressant versus placebo response: Evidence from the EMBARC study. Psychol. Med. 49, 1118–1127 (2019).

39. Bjork, J. M., Straub, L. K., Provost, R. G. & Neale, M. C. The ABCD study of neurodevelopment: Identifying neurocircuit targets for prevention and treatment of adolescent substance abuse. Curr. Treat. Options Psychiatry 4, 196–209 (2017).

40. Human Connectome Project. HCP-Aging Lifespan 2.0 Release - Connectome. https://www.humanconnectome.org/study/hcp-lifespan-aging/document/hcp-aging-20-release.

41. Kennedy, K. M. et al. Age trajectories of functional activation under conditions of low and high processing demands: an adult lifespan fMRI study of the aging brain. Neuroimage 104, 21–34 (2015).

42. Fan, L., Klein, H., Bass, E., Springfield, C. & Pinkham, A. Amygdala Hyperconnectivity in the Paranoid State: A Transdiagnostic Study. J. Psychiatr. Res. 138, 117–124 (2021).

43. Addington, J. et al. North American Prodrome Longitudinal Study (NAPLS 3): Methods and baseline description. Schizophr Res 243, 262–267 (2022).

44. Lenze, E. J., Voineskos, A. N., Butters, M. A. & Karp, J. F. Stopping Cognitive Decline in Patients With Late-Life Depression: A New Front in the Fight Against Dementia. Am. J. Geriatr. Psychiatry Off. J. Am. Assoc. Geriatr. Psychiatry 26, 828–834 (2018).

45. Kohli, J. S., Kinnear, M. K., Martindale, I. A., Carper, R. A. & Müller, R.-A. Regionally decreased gyrification in middle-aged adults with autism spectrum disorders. Neurology 93, e1900–e1905 (2019).

46. Ciarleglio, A. J. et al. A predictive model for conversion to psychosis in clinical high-risk patients. Psychol. Med. 49, 1128–1137 (2019).

47. Human Connectome Project. Perturbation of the Treatment of Resistant Depression Connectome by Fast-Acting Therapies (PDC) - Connectome. https://www.humanconnectome.org/study/crhd-perturbation-treatment-resistant-depression-connectome-fast-acting-therapies/data-releases.

48. Geeraert, B. L., Chamberland, M., Lebel, R. M. & Lebel, C. Multimodal principal component analysis to identify major features of white matter structure and links to reading. PLOS ONE 15, e0233244 (2020).

49. Human Connectome Project. HCP-Development Lifespan 2.0 Release - Connectome. https://www.humanconnectome.org/study/hcp-lifespan-development/document/hcp-development-20-release.

50. Human Connectome Project. BANDA Release 1.1 - Connectome. https://www.humanconnectome.org/study/connectomes-related-anxiety-depression/document/banda-release-11.

51. Horta, M., Polk, R. & Ebner, N. Single Dose Intranasal Oxytocin Administration: Data from Healthy Younger and Older Adults. Openneuro 10.18112/OPENNEURO.DS004725.V1.0.1 (2023).

52. Human Connectome Project. 1200 Subjects Data Release - Connectome. https://www.humanconnectome.org/study/hcp-young-adult/document/1200-subjects-data-release.

53. Binder, J., et al. Epilepsy Connectome Project. https://osf.io/exbt4 (2023).

54. RePORT ⟩ RePORTER. https://reporter.nih.gov/project-details/9748600.

55. Wannan, C. M. J. et al. Accelerating Medicines Partnership® Schizophrenia (AMP® SCZ): Rationale and Study Design of the Largest Global Prospective Cohort Study of Clinical High Risk for Psychosis. Schizophr. Bull. 50, 496–512 (2024).

56. Human Connectome Project. Human Connectome Project for Early Psychosis - Connectome. https://www.humanconnectome.org/study/human-connectome-project-for-early-psychosis/data-releases.

57. Seok, D. et al. Dimensional connectomics of anxious misery, a human connectome study related to human disease: Overview of protocol and data quality. NeuroImage Clin. 28, 102489 (2020).

58. Diniz, B. S. et al. Mild cognitive impairment and major depressive disorder are associated with molecular senescence abnormalities in older adults. Alzheimers Dement. Transl. Res. Clin. Interv. 7, e12129 (2021).

59. Meram, T. D. et al. Evoking network profiles of the dorsal anterior cingulate in youth with Obsessive-Compulsive Disorder during motor control and working memory. J. Psychiatr. Res. 132, 72–83 (2021).

60. Baran, B. et al. Sleep and wake markers of thalamocortical functioning in early-course psychosis and first-degree relatives. Schizophrenia 12, 40 (2026).

